# Barthelonids represent a deep-branching Metamonad clade with mitochondrion-related organelles generating no ATP

**DOI:** 10.1101/805762

**Authors:** Euki Yazaki, Keitaro Kume, Takashi Shiratori, Yana Eglit, Goro Tanifuji, Ryo Harada, Alastair G.B. Simpson, Ken-ichiro Ishida, Tetsuo Hashimoto, Yuji Inagaki

## Abstract

We here report the phylogenetic position of barthelonids, small anaerobic flagellates previously examined using light microscopy alone. *Barthelona* spp. were isolated from geographically distinct regions and we established five laboratory strains. Transcriptomic data generated from one *Barthelona* strain (PAP020) was used for large-scale, multi-gene phylogenetic (phylogenomic) analyses. Our analyses robustly placed strain PAP020 at the base of the Fornicata clade, indicating that barthelonids represent a deep-branching Metamonad clade. Considering the anaerobic/microaerophilic nature of barthelonids and preliminary electron microscopy observations on strain PAP020, we suspected that barthelonids possess functionally and structurally reduced mitochondria (i.e. mitochondrion-related organelles or MROs). The metabolic pathways localized in the MRO of strain PAP020 were predicted based on its transcriptomic data and compared with those in the MROs of fornicates. Strain PAP020 is most likely incapable of generating ATP in the MRO, as no mitochondrial/MRO enzymes involved in substrate-level phosphorylation were detected. Instead, we detected the putative cytosolic ATP-generating enzyme (acetyl-CoA synthetase), suggesting that strain PAP020 depends on ATP generated in the cytosol. We propose two separate losses of substrate-level phosphorylation from the MRO in the clade containing barthelonids and (other) fornicates.

## Introduction

Elucidating the evolutionary relationships among the major groups of eukaryotes is one of the most fundamental but unsettled questions in biology. It is widely accepted that large-scale molecular data for phylogenetic analyses (so-called phylogenomic data) are indispensable to infer ancient splits in the tree of eukaryotes (Burki 2014; Burki et al. 2019; Keeling and Burki 2019). Preparing phylogenomic data has been greatly advanced by the recent technological improvements in sequencing that generate a large amount of molecular data at an affordable cost and in a reasonable time-frame (Bleidorn 2016; Vincent et al. 2017). Further, some recent phylogenomic analyses have included uncultured microbial eukaryotes (e.g., Lax et al. 2018), since the libraries for sequencing of the whole-genome/transcriptome can be prepared from a small number of cells (or even a single cell) isolated from an environment sample (Kolisko et al. 2014; Strassert et al. 2019).

Despite these advances in experimental techniques, it is realistic to assume that no current phylogenomic analysis has covered the true diversity of eukaryotes. A large number of extant microbial eukaryotes have never been examined using transcriptomic or genomic techniques, and some of them may hold the keys to resolving important unanswered questions in eukaryotic phylogeny and evolution. Thus, to reconstruct the evolutionary relationships among the major eukaryotic assemblages to a resolution that is both accurate and informative, the taxon sampling in phylogenomic analyses has been improved by targeting two classes of organisms: (i) Novel microbial eukaryotes that represent lineages that were previously unknown to science, and (ii) “orphan eukaryotes” that had been reported before, but whose evolutionary affiliations were unresolved by morphological examinations and/or single-gene phylogenies (Zhao et al. 2012; Kamikawa et al. 2014; Yabuki et al. 2014; Burki et al. 2016; Janouškovec et al. 2017; Brown et al. 2018; Lax et al. 2018; Gawryluk et al. 2019; Strassert et al. 2019).

Many of “orphan eukaryotes” were described based solely on morphological information prior to the regular use of gene sequences in phylogenetic/taxonomic studies. One such organism is the small free-living heterotrophic biflagellate *Barthelona vulgaris* (Bernard et al. 2000). The initial description of *B. vulgaris* was based on light microscopy observations of cells isolated from marine sediment from Quibray Bay, Australia, and maintained temporarily in nominally anoxic crude culture (Bernard et al. 2000). The morphospecies was later identified at different geographical locations (Lee 2002; Lee 2006) but never examined with methods incorporating molecular data. These past studies identified no special morphological similarity between *B. vulgaris* and any eukaryotes described to date at the morphological level (Bernard et al. 2000; Lee 2002; Lee 2006). Thus, to clarify the phylogenetic placement of *B. vulgaris* in the tree of eukaryotes, molecular phylogenetic analyses are required, preferably at the “phylogenomic” scale.

We here report five laboratory strains of *Barthelona* (EYP1702, FB11, LRM2, PAP020 and PCE; Figs. 1A-E) isolated from separate geographical regions, and infer their phylogenetic positions assessed by analyzing both SSU rDNA and phylogenomic data. A SSU rDNA phylogeny robustly united all of the *Barthelona* strains together, but the precise placement of *Barthelona* spp. among other eukaryotes remained inconclusive. To infer the precise phylogenetic position of barthelonids, we obtained a transcriptome data from strain PAP020, and analyzed its phylogenetic position from a eukaryote-wide dataset containing 148 genes. The transcriptome data of strain PAP020 was also used for reconstructing the metabolic pathways in a functionally and structurally reduced mitochondrion that is the result of adaptation to anaerobiosis.

**Figure 1.**
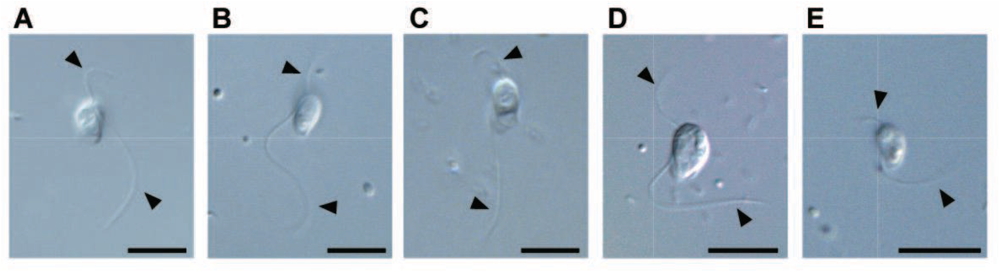
Light micrographs of *Barthelona* spp. studied in this study. Strains PAP020, FB11, LRM2, EYP1702 and PCE are shown in (A), (B), (C), (D), and (E), respectively. Flagella are marked by arrowheads. Scale bars = 10 μm.

## Results and Discussion

### Phylogenetic position of barthelonids

Overall, the maximum-likelihood (ML) and Bayesian phylogenetic analyses of small subunit ribosomal RNA gene (SSU rDNA) sequences resolved known major eukaryote groups with moderate to high statistical support values, but the backbone of the tree remained unresolved (Fig. 2). In the SSU rDNA tree, all of *Barthelona* sp. strains PAP020, EYP1702, FB11, PCE, and LRM2 grouped together with a ML bootstrap value (MLBP) of 83% and a Bayesian posterior probability (BPP) of 0.98. In this *Barthelona* clade, strains EYP1702 and PCE were the earliest and second earliest diverging taxa, respectively, and strains PAP020, LRM2 and FB11 formed a tight subclade. The *Barthelona* clade was sister to a Fornicata clade comprising *Carpediemonas membranifera, Kipferlia bialata, Dysnectes brevis, Retortamonas* sp. and *Giardia intestinalis* (Fig. 2), but statistical support was equivocal (MLBP 56%; BPP 0.86). This possible affinity between *Barthelona* and fornicates in the SSU rDNA phylogeny is provocative, as both lineages thrive in oxygen-poor environments and possess double-membrane bound MROs instead of typical mitochondria (Simpson and Patterson 1999; Tovar et al. 2003; Yubuki et al. 2007; Yubuki et al. 2013; Kulda et al. 2017; see Fig. S2 for the putative MRO in strain PAP020). Thus, we took a phylogenomic approach to more robustly resolve the position of barthelonids in the tree of eukaryotes.

**Figure 2.**
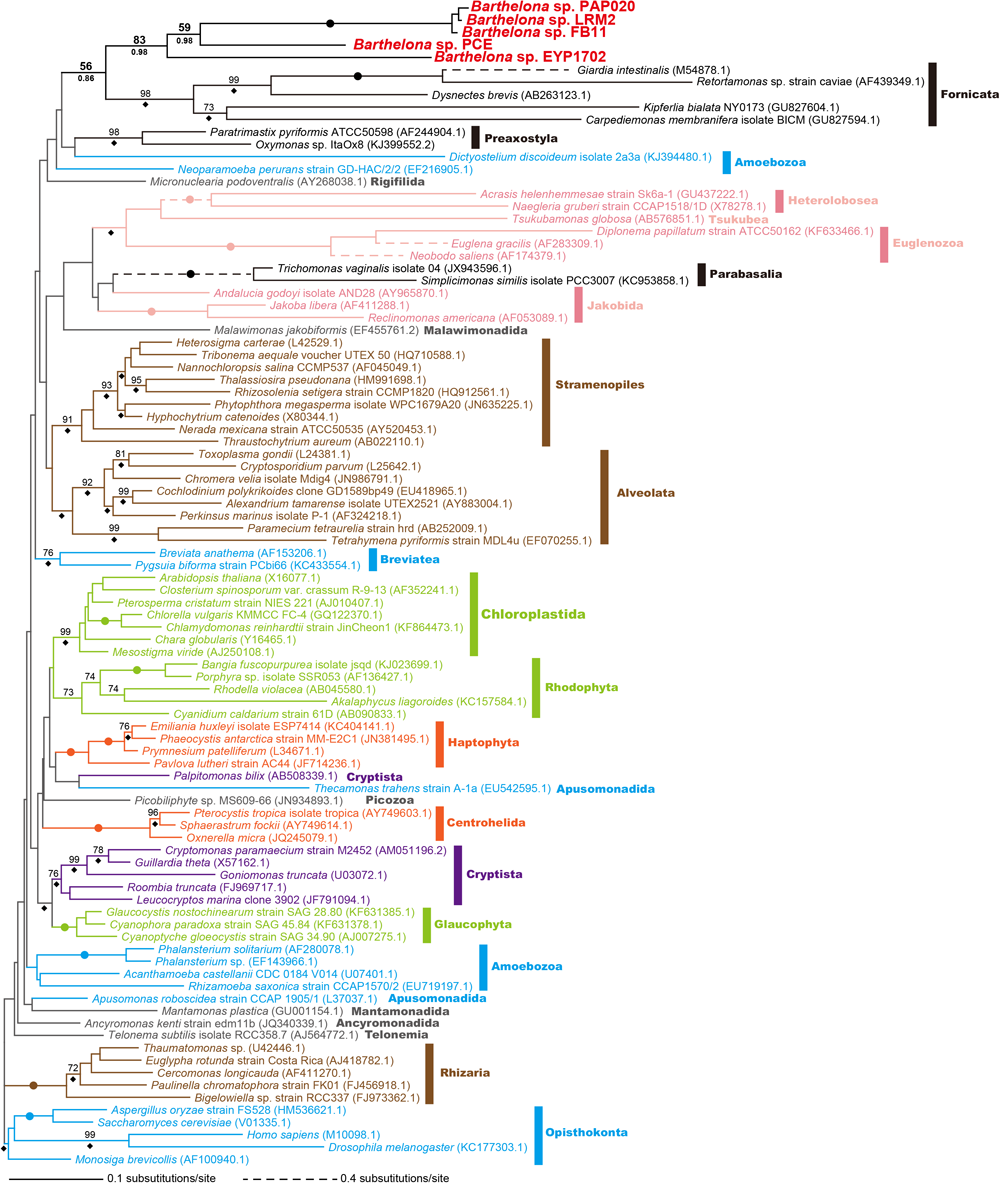
Global eukaryotic phylogeny inferred from small subunit ribosomal DNA sequences. The tree topology was inferred using the maximum-likelihood (ML) method and ML bootstrap values (MLBPs) and Bayesian posterior probabilities (BPPs) were mapped on the ML tree. The nodes marked by dots were supported by MLBPs of 100% and BPPs of 1.0. MLBPs less than 70% are not shown. BPPs of 0.95 or more are marked by diamonds.

As anticipated, both ML and Bayesian phylogenetic analyses of a multi-gene alignment comprising 148 genes (148-gene alignment) provided us deeper insights into the backbone of the tree of eukaryotes (Fig. 3) than the SSU rDNA analyses (Fig. 2). The backbone tree topology and statistical support values (Fig. 3) agreed largely with those reported in Kamikawa et al. (2014), Yabuki et al. (2014) and Yabuki et al. (2018), which analyzed multi-gene alignments generated from the same core set of 157 single-gene alignments with mostly similar taxon sampling. The topology includes well established clades including SAR, Amorphea, Cryptophyceae and Discoba, but, as is common, did not infer a monophyletic Archaeplastida (Cenci et al. 2018; Strassert et al. 2019). Likewise, the 148-gene phylogeny recovered neither the clade of *Telonema subtilis* and SAR (“T-SAR”; Strassert et al. 2019) nor that of centrohelids and haptophytes (Haptista; Burki et al. 2016). We suspect that large proportions of missing data in the sequence of *T. subtilis* and the single included centrohelid (34 and 35%, respectively), which derived from the transcriptomic data generated by 454 pyrosequencing (Burki et al. 2009), hindered the recoveries of T-SAR and Haptista in the 148-gene phylogeny.

**Figure 3.**
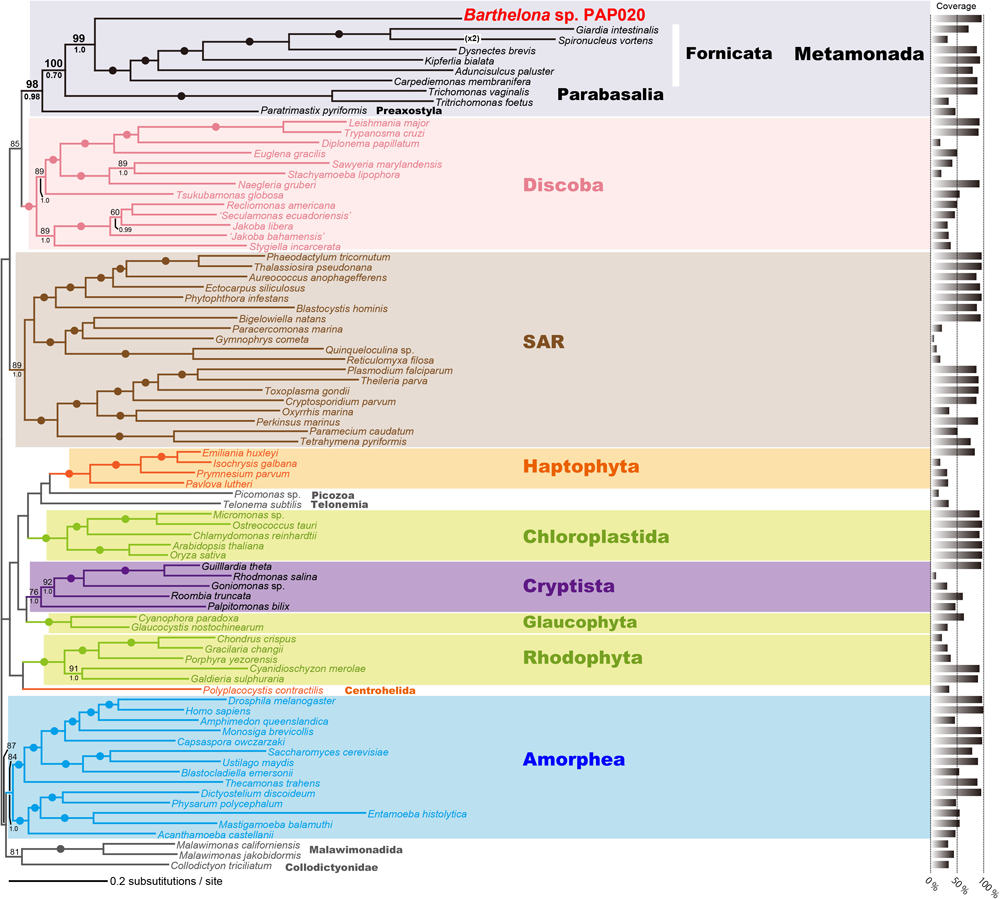
Global eukaryotic phylogeny inferred from a 148-gene alignment. The tree topology was inferred using the maximum-likelihood (ML) method; ML bootstrap values (MLBPs) and Bayesian posterior probabilities (BPPs) were mapped on the ML tree. The Bayesian analysis recovered and identical overall topology. The nodes marked by dots were supported by MLBPs of 98% or more, and BPPs of 0.95 or more. MLBPs less than 60% or BPPs below 0.80 are not shown. The bar graph for each taxon indicates the percent coverage of the amino acid positions in the 148-gene analyses.

The 148-gene phylogeny grouped *Barthelona* sp. strain PAP020 and 6 fornicates together with a MLBP of 99% and a BPP of 1.0 (Fig. 3). In this clade, strain PAP020 occupied the basal position, which was supported fully by both ML and Bayesian analyses. The clade of strain PAP020 and fornicates was connected sequentially with parabasalids (MLBP 100%; BPP 0.70), then with *Paratrimastix pyriformis* (representing Preaxostyla), to form the Metamonada clade with a MLBP of 98% and a BPP of 0.98 (Fig. 3). Support for these relationships was hardly affected by exclusion of rapidly evolving alignment positions, until >60% of site were excluded (Fig. S1). We applied the ML tree and three alternative trees, wherein strain PAP020 branched at the base of the Parabasalia clade, the clade of Fornicata + Parabasalia and the Metamonada clade, to an approximately unbiased (AU) test, and all of the alternative trees were rejected (p < 0.001). The results from the phylogenetic analyses of 148-gene alignment consistently and robustly indicated that barthelonids are a previously overlooked Metamonada lineage, which has a specific affinity with the Fornicata clade.

There are two uncertain issues related to the taxonomic treatment of barthelonids for future studies. Firstly, molecular phylogenetic analyses alone cannot determine whether barthelonids are (a) a sister taxon to Fornicata, or (b) the deepest known branch within the taxon of Fornicata. Fornicata is defined by a key ultrastructural characteristic in the flagellar apparatus, namely the so-called “B fiber” forms a conspicuous arching bridge between the two flagellar roots supporting the ventral feeding groove (Simpson 2003). Therefore, we need to investigate the ultrastructure of barthelonid cells in detail for their higher-level taxonomic treatment. The second issue for future studies is whether it is appropriate to classify all of the five strains examined in this study into a single genus *Barthelona*. In the *Barthelona* clade recovered in the SSU rDNA phylogeny (Fig. 2), strains PAP020, LRM2 and FB11 appeared to be closely related to one another but are distant from strains PCE and EYP1702. We need to assess their morphological characteristics to settle this issue.

### Lack of substrate-level phosphorylation in the mitochondrion-related organelle of *Barthelona* sp. PAP020

All of the *Barthelona* strains assessed in this study (strains PAP020, EYP1702, PCE, LRM2 and FB11) are grown under oxygen-poor conditions in the laboratory. Our preliminary ultrastructural observation of strain PAP020 did not reveal a typical mitochondrion. Instead we observed a densely stained, double membrane-bounded organelle (Fig. S2). As all metamonads studied so far lack typical mitochondria, we suspect that the double membrane-bounded organelle identified in strain PAP020 is the MRO. According to the phylogenetic position of barthelonids deduced from the SSU rDNA and 148-gene phylogeny (Figs. 2 & 3), the metabolic pathways retained in the barthelonid MROs are significant to infer the evolutionary history of the MROs in the Fornicata clade.

Leger et al. (2017) proposed that the ancestral fornicate species possessed an MRO with a metabolic capacity similar to that of the hydrogenosomes in parabasalids like *Trichomonas vaginalis*. Thus, we surveyed the transcriptomic data from strain PAP020 for transcripts encoding hydrogenosomal/MRO proteins that are homologous to *Trichomonas* proteins localized in the hydrogenosome. Strain PAP020 was predicted to possess the MRO proteins involved in hydrogen production, pyruvate metabolism, amino acid metabolism, FeS cluster assembly, anti-oxidant system and protein modification (chaperones and proteases) (Figure 4A; purple and grey ellipses represent the proteins found and not found, respectively; see also Table S2). This suggests that the overall function of the MRO of strain PAP020 is similar to that of the *Trichomonas* hydrogenosome, except in ATP generation capacity. We did not identify any transcripts encoding two enzymes for anaerobic ATP generation through substrate-level phosphorylation, namely (i) acetate:succinate CoA transferase (ASCT) that transfers coenzyme A (CoA) from acetyl-CoA to succinate and (ii) succinyl-CoA synthase (SCS) that phosphorylates ADP to produce ATP coupled with converting succinyl-CoA back to succinate. We propose that strain PAP020 genuinely lacks ASCT and SCS and that its MRO is incapable of generating ATP.

**Figure 4.**
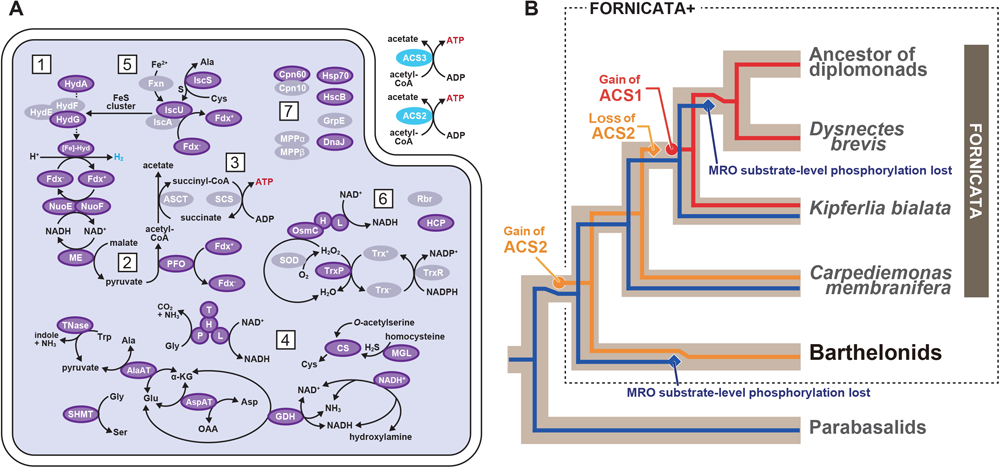
Function and evolution of the mitochondrion-related organelles (MRO) of *Barthelona* sp. strain PAP020. A. Reconstructed metabolic pathways in the MRO of strain PAP020. Dark purple ellipses indicate that the transcripts encoding hydrogenosomal/MRO proteins were detected in the *Barthelona* RNA-seq data, and their N-termini were predicted as transit peptides for mitochondria/MRO by MitoFates (Fukasawa et al. 2015) and/or NommPred (Kume et al. 2018). Pale purple ellipses indicate putative hydrogenosomal/MRO proteins lacking N-terminal sequence information or those with N-terminal extensions that were not predicted as mitochondria/MRO localizing by MitoFates and NommPred. Hydrogenosomal/MRO proteins shown in grey ellipses represent the absence of the corresponding transcripts in the *Barthelona* RNA-seq data. Strain PAP020 possesses two acetatyl CoA synthases (ACS), one corresponds to the cytosolic ACS of multiple fornicates (ACS2) and the other showed no clear phylogenetic affinity to any known ACS (ACS3; see Fig. S3). We regard ACS2 as a cytosolic protein in strain PAP020 (blue ellipse). As there is no hint for the subcellular localization of ACS3, this version is omitted from the figure. 1, H2-synthesis; 2, pyruvate metabolism; 3, substrate-level phosphorylation; 4, amino acid metabolism; 5, Fe-S cluster assembly; 6, anti-oxidant system; 7, protein modification. Abbreviations: Ala, alanine; Asp, aspartic acid; Cys, cysteine; Glu, glutamic acid; Gly, glycine; α-KG, α-ketoglutaric acid; Trp, tryptophan; OAA, oxaloacetic acid; NAD^+^/NADH, nicotinamide adenine dinucleotide; NADP^+^/NADPH, nicotinamide adenine dinucleotide phosphate; HydE/F/G, hydrogenase maturases E/F/G; HydA/[Fe]-Hyd, hydrogenase; Fdx, ferredoxin; NuoE/F, 24/51 kDa of mitochondrial NADH:ubiquinone oxidoreductase; ME, malic enzyme; PFO, pyruvate:ferredoxin oxidoreductase; ASCT, acetate:succinyl-CoA transferase; H/L/P/T, glycine cleavage system protein H/L/P/T; AlaAT, alanine aminotransferase; AspAT, aspartate aminotransferase; TNase, tryptophanse; GDH, glutamate dehydrogenase; HCP, hybrid-cluster protein; SHMT, serine hydroxymethyltransferase; CS, cysteine synthase; MGL, monoacylglycerol lipase; Fxn,; IscS, cysteine desulfurase; IscU/A, iron-sulfur cluster assembly protein; Fxn, frataxin; OsmC, osmotically inducible protein; SOD, superoxide dismutase; Trx, thioredoxin; TrxR, thioredoxin reductases; TrxP, thioredoxin peroxidase; Rbr, rubrerythrin; Cpn60/10, chaperonin 60/10; Hsp70, heat shock protein 70; HscB, heat shock cognate B; GrpE, nucleotide exchange factor for DnaK; DnaJ, heat shock protein 40; MPPα/β, mitochondrial processing peptidase α/β. B. Evolution of ATP generation in barthelonids, parabasalids and selected fornicates. In the clade of fornicates and barthelonids (Fornicata+ clade), substrate-level phosphorylation (blue) was lost on two separate branches. The cytosolic ACS2 (yellow), which was established at the base of the Fornicata+ clade, was replaced by an evolutionarily distinct type of ACS (ACS1; red) during the evolution of fornicates.

We additionally surveyed the PAP020 data for transcripts encoding cytosol-localizing acetyl-CoA synthase (ACS), which is an alternative mechanism to generate ATP in fornicate cells. Intriguingly, two distinct ACS sequences were retrieved, designated here as ACS2 and ACS3. Although the transcripts encoding both ACS versions most likely cover their N-termini, neither of them was predicted to bear the typical signal to be localized in mitochondria or MROs (i.e. an inferred N-terminal transit peptide). The abundances of the ACS2 and ACS3 transcripts in strain PAP020 were 2249 and 2208 Transcripts Per kilobase Million (TPM; Li and Dewey 2011), respectively, implying that the two *Barthelona* ACS genes are indistinguishable at the transcription level. We subjected the two ACS sequences to a phylogenetic analysis along with the homologues sampled from diverse bacteria, archaea and eukaryotes (Fig S3). The PAP020 ACS2 sequence formed a clade with fornicate “ACS2” sequences, which Leger et al. (2017) proposed to be cytosolic enzymes. Thus, we suggest that ACS2 is most likely a cytosolic enzyme in strain PAP020 as well. The ACS phylogeny recovered no strong affinity between PAP020 ACS3 sequence and other homologues (Fig. S3). Neither of our analyses on the ACS3 sequence provided any positive support for MRO localization, and we tentatively consider ACS3 as a cytosolic enzyme in strain PAP020. Altogether, we conclude that strain PAP020 depends entirely on ATP generated by the two cytosol-localizing ACS, as its MRO lacks substrate-level phosphorylation. For a better understanding of ATP synthesis in this organism, the precise subcellular localizations of ACS2 and ACS3 in strain PAP020 need to be confirmed experimentally in the future.

Leger et al. (2017) proposed a complex evolution of ATP-generating mechanisms in the Fornicata clade, as follows: (i) The ancestral fornicate species possessed both substrate-level phosphorylation in the MRO as well as ACS2 in the cytosol. (ii) Substrate-level phosphorylation has been inherited vertically to the extant fornicate species, except *D. brevis* and diplomonads (see below). (iii) During the evolution of Fornicata, the ancestral cytosol-localizing ACS (i.e. ACS2) was replaced by an evolutionarily distinct ACS (ACS1). (iv) The redundancy in the ATP-generating system allowed the secondary loss of substrate-level phosphorylation in the MRO prior to the separation of the *D. brevis* plus diplomonad clade. We here extend the scenario proposed by Leger et al. (2017) by incorporating the data from *Barthelona* sp. strain PAP020 (See Fig. 4B). Acquisition of ACS2 was hypothesized at the base of the Fornicata clade in the previous work (Leger et al. 2017), but after assessing the data from stain PAP020, this particular event needs to be pushed back at least to the common ancestor of fornicates and barthelonids, as strain PAP020 and multiple early-branching CLOs (e.g., *C. membranifera)* share ACS2. It is worthy to note that acquisition of ACS2 may extend back to the last common metamonad ancestor, since a possibly directly related ACS2 is also present in *Trimastix* (Fig. S3). Secondly, as barthelonids are distantly related to *D. brevis* and diplomonads, loss of substrate-level phosphorylation in barthelonid MROs can be assumed to have occurred independently from the loss in the common ancestor of *D. brevis* and diplomonads (highlighted by blue diamonds in Fig. 3B). Further, barthelonids and the common ancestor of *D. brevis* and diplomonads seem to have accommodated the loss of MRO-localized substrate-level phosphorylation via possessing evolutionarily distinct ACS homologs (ACS2 and ACS1, represented by yellow and red lines, respectively in Fig. 4B).

In the current study, we reconstructed the metabolic pathways in the MRO of only one of the five strains of *Barthelona* sp. We anticipate that stains PAP020, LRM2 and FB11, which formed a tight clade in the SSU rDNA phylogeny (Fig. 2), may have MROs with the same or a very similar set of metabolic pathways. In future studies, it is important to reconstruct the metabolic pathways in the MROs of strains PCE and/or EYP1702 to further resolve the evolution of MROs and anaerobic metabolism in the Metamonada clade. Considering the large evolutionary distance between PCE/EYP1702 and PAP020/LRM2/FB11 in the SSU rDNA phylogeny (Fig. 2), we may find that the MRO functions of strains PCE and EYP1702 are substantially different from that of strain PAP020 deduced in the current study.

## Materials & Methods

### Isolation and Cultivation

We established five laboratory strains of *Barthelona* sp. in this study (Fig. 1A-E). Strains PAP020 and EYP1702 (Figs. 1A & 1D) were isolated from anaerobic mangrove sediments collected at a seawater lake in the Republic of Palau in November 2011 and October 2017, respectively. The laboratory cultures have been maintained in mTYGM-9 medium (http://mcc.nies.go.jp/medium/ja/mtygm9.pdf) with prey bacteria at 18-20 °C. An anaerobic environment within the laboratory cultures was created by the respiration of prey bacteria. LRM2 (Fig 1B) was isolated from mud of a defunct saltern (now normal salinity) on the Ebre Delta near San Carles de la Ràpita, Catalonia, Spain in February 2015. FB11 (Fig1C) was isolated from False Bay, an intertidal mud flat on San Juan Island, WA, US in June 2015. PCE (Fig 1E) was isolated from intertidal sediment near Cavendish, PEI, Canada in July, 2016. The established cultures were maintained with co-cultured bacteria on 3% LB in sterile natural seawater at 18-21°C.

### SSU rDNA phylogenetic analysis

Total DNA samples of *Barthelona* sp. strains PAP020, EYP1702, FB11, PCE and LRM2 were extracted from the cultured cells using a DNeasy Plant mini kit (Qiagen) or NucleoSpin® Tissue kit (Macherey-Nagel). Near-complete SSU rDNA fragments were amplified from each DNA sample by PCR, using either primers SR1 and SR12 (Nakayama et al. 1998) or 18F and 18R (Yabuki et al. 2010). The amplification program consisted of 30 cycles of denaturation at 94 °C for 30 s, annealing at 55 °C for 30 s and extension at 72 °C for 90 s. The amplified product was gel-purified, cloned and sequenced by the Sanger method.

We aligned the SSU rDNA sequences of the five *Barthelona* strains with those of 91 phylogenetically diverse eukaryotes by using MAFFT 7.205 (Katoh 2002; Katoh and Standley 2014). After manual exclusion of ambiguously aligned positions, 1,573 nucleotide positions were subjected to ML phylogenetic analyses by using IQTREE v 1. 5. 4 (Nguyen et al. 2015) with the GTR + R6 model, with MLBPs derived from 500 non-parametric bootstrap replicates. The SSU rDNA alignment was also subjected to Bayesian phylogenetic analysis using MrBayes 3.2.3 (Ronquist et al. 2012) with GTR + Γ model. The Markov Chain Monte Carlo (MCMC) run was performed with one cold and three heated chains with default chain temperatures. We ran 3,000,000 generations, and sampled log-likelihood scores and trees with branch lengths every 1,000 generations (the stationarity was confirmed by plotting the log-likelihoods sampled during the MCMC). The first 25% generations were discarded as burn-in. The consensus tree with branch lengths and BPPs were calculated from the remaining trees.

### RNA-seq analyses

We conducted two RNA-seq runs of *Barthelona* sp. strain PAP020. The sequence reads from the first analysis was used for a phylogenomic analysis assessing the position of *Barthelona* spp. in the tree of eukaryotes, while those from the second sequencing run were for surveying the proteins localized in the mitochondrial-related organelle (MRO) in strain PAP020 (see below).

For the first RNA-seq run, PAP020 cells, together with bacterial cells in the culture medium, were harvested and subjected to RNA extraction using TRIzol (Life Technologies) by following the manufacturer’s protocol. We shipped the RNA sample to a biotech company (Hokkaido System Science) for cDNA library construction and subsequent sequencing using the Illumina HiSeq 2500 platform, which generated 2.9 x 10^7^ paired-end 100-bp reads (2.9 Gb in total). The initial reads were then assembled into 29,251 unique contigs by TRINITY (Grabherr et al. 2011; Haas et al. 2013).

For the second RNA-seq run, we separated PAP020 cells from the bacterial cells in the culture medium by a gradient centrifugation using Optiprep (Axis Shield), as reported previously (Tanifuji et al. 2018), with slight modifications (the Optiprep solution containing the eukaryotic cells and bacteria was centrifuged at 2,000 *g* for 20 min, instead of 800 *g* for 20 min). Total RNA was extracted from the harvested eukaryote-enriched fraction, using TRIzol by following the manufacturer’s protocol. Poly-A tailed RNAs in the RNA sample described above were purified with a Dynabeads™ mRNA Purification Kit (Thermo Fisher Scientific), and then used to construct the cDNA library using the SMART-Seq v4 Ultra Low Input RNA Kit (Takara Bio USA) for Sequencing and Nextera XT DNA Library Preparation Kit (Illumina). The resultant cDNA library was sequenced with the Illumina Miseq platform, yielding 3.7 x 10^7^ paired-end 300-bp sequence reads (8.6 Gb in total). These were assembled into 21,286 unique contigs using TRINITY.

### Phylogenomic analyses

To elucidate the phylogenetic position of *Barthelona* sp. strain PAP020, we prepared a phylogenomic alignment by updating an existing dataset comprising 157 genes (Kamikawa et al. 2014; Yabuki et al. 2014; Yabuki et al. 2018, see Table S1). For each of these 157 genes, we added the homologous sequences retrieved from the transcriptomic data of strain PAP020 (this study) and four fornicates (*Carpediemonas membranifera, Aduncisulcus paluster, Kipferlia bialata* and *Dysnectes brevis*; Leger et al. 2017). Each single-gene alignment was aligned individually by MAFFT 7.205 with the L-INS-i algorithm followed by manual correction and exclusion of ambiguously aligned positions. For each of the single-gene alignments, the ML phylogenetic tree was inferred by RAxML 8.1.20 (Stamatakis 2014) under the LG + Γ + F model with robustness assessed with a 100 replicate bootstrap analysis.

Individual single-gene trees were inspected to identify the alignments bearing aberrant phylogenetic signal that disagreed strongly with any of a set of well-established monophyletic assemblages in the tree of eukaryotes, namely Opisthokonta, Amoebozoa, Alveolata, Stramenopiles, Rhizaria, Rhodophyta, Viridiplantae, Glaucophyta, Haptophyta, Cryptophyta, Jakobida, Euglenozoa, Heterolobosea, Diplomonadida, Parabasalia and Malawimonadidae. Nine out of the 157 single-gene alignments were found to bear idiosyncratic phylogenetic signal and were excluded from the phylogenomic analyses described below. After inspection of single-gene alignments/trees, the remaining 148 single-gene alignments (Table S1) were concatenated into a single phylogenomic alignment containing 83 taxa with 38,816 unambiguously aligned amino acid positions (148-gene alignment). The coverage for each single-gene alignment is summarized in Table S1.

ML analyses of 148-gene alignment were conducted by using IQ-TREE v. 1.5.4 with the LG + Γ + F + C60 + PMSF (posterior mean site frequencies) model (Wang et al. 2018) and robustness evaluated with a ML bootstrap analysis on 100 replicates. We also conducted a Bayesian phylogenetic analysis with the CAT + GTR model using PHYLOBAYES 1.5a (Lartillot and Philippe 2004; Lartillot and Philippe 2006; Lartillot et al. 2007). In each analysis, two MCMC runs were run for 5,000 cycles with “burn-in” of 1,250 (‘maxdiff value was 0.96743). The consensus tree with branch lengths and Bayesian posterior probabilities (BPPs) were calculated from the remaining trees.

The phylogenetic position of *Barthelona* sp. strain PAP020 inferred from the 148-gene alignment was assessed by an approximately unbiased test (Shimodaira 2002). We modified the ML tree to prepare four alternative tree topologies, in which strain PAP020 branches 1) at the base of the Parabasalia clade, 2) at the base of the clade of parabasalids and fornicates, 3) with *Paratrimastix pyriformis*, and 4) at the base of the Metamonada clade. Site likelihood data were calculated over each of the five trees examined (ML plus four alternative trees) using IQ-TREE and then analyzed in CONSEL ver.0.20 (Shimodaira and Hasegawa 2001) with the default settings.

We evaluated the contribution of fast-evolving sites in the 148-gene alignment to the position of *Barthelona* sp. strain PAP020. Individual rates for sites were calculated over the ML tree topology using DIST_EST (Susko et al. 2003) with the LG + Γ + F model. Fast-evolving sites were progressively removed from the original 148-gene alignment in 4,000-position increments, and each of the resulting alignments was subjected to 100 replicate rapid ML bootstrap analysis with RAxML 8.1.20 with the LG + Γ + F model.

### Prediction of proteins localized in the mitochondrion-related organelle in *Barthelona* sp. PAP020

We searched for mRNA sequences encoding proteins predicted to be localized to the mitochondrion-related organelle (MRO) in *Barthelona* sp. strain PAP020, as well as those involved in anaerobic ATP generation. For this we searched among the contigs generated from the second RNA-seq experiment by TBLASTN, using the hydrogenosomal/MRO proteins in *Trichomonas vaginalis* and *Giardia intestinalis* as the queries (Leger et al. 2017). The amino acid sequences deduced from the contigs retrieved by the first BLAST searches were then subjected to BLASTP analyses against NCBI nr database to exclude false positives. The domain structures of the putative MRO proteins were examined using hmmscan 3.1 (http://hmmer.org). We inspected each of the putative MRO proteins for potential mitochondrial targeting sequences using MitoFates (Fukasawa et al. 2015) with default parameters for the fungal sequences, and NommPred (Kume et al. 2018) with parameters for canonical mitochondria and MRO.

## Acknowledgements

This work was supported in part by a fund from the grants from the Japan Society for the Promotion of Science (18KK0203 and 19H03280 awarded to YI; 15H05231 and 19KK0185 to TH) and by the “Tree of Life” research project of University of Tsukuba, as well as Natural Sciences and Engineering Research Council (Canada) Discovery (Grant 298366-2014 to AGBS). We thank Dr. Noèlia Carrasco for providing access to the Delta de l’Ebre sampling location for strain LRM2. The SSU rDNA sequences of *Barthelona* spp. were deposited in GenBank/EMBL/DDBJ database under the accession nos. LC506386–LC506390. The transcriptome data of strain PAP020 were deposited in DDBJ Sequence Archive under the accession nos. SRA### and SRA$$$$.

---

**Table S1: Amino acid positions and coverages of the 148 single-gene alignments used in this study.**

**Table S2: Putative MRO proteins of *Barthelona* sp. strain PAP020 and their predicted subcellular localizations.**

**Figure S1:**
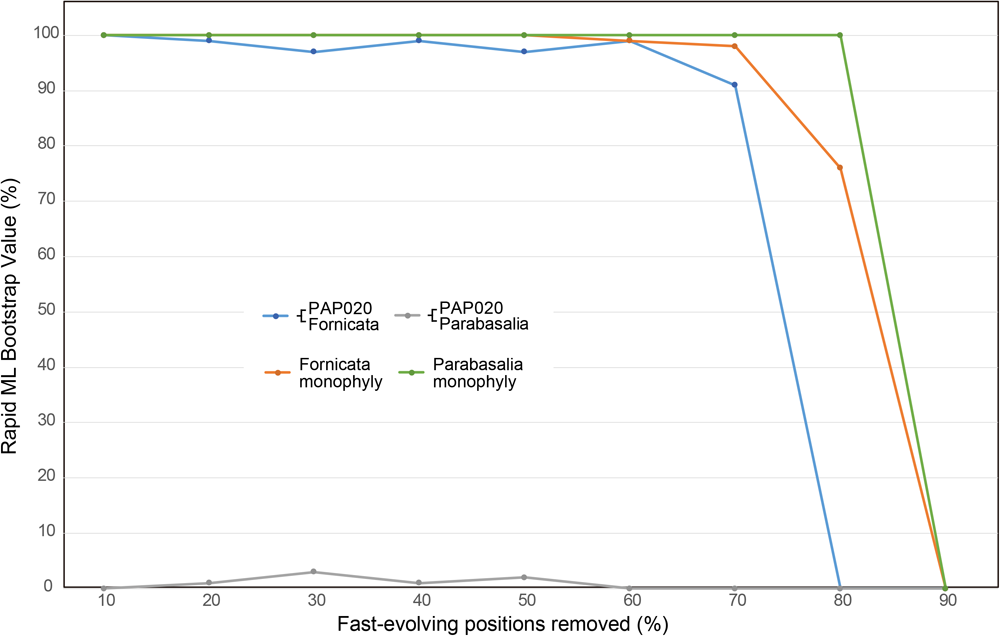
The impact of removal of fast-evolving alignment positions on the phylogenetic relationship among fornicates, parabasalids and *Barthelona* sp. strain PAP020. Fast-evolving positions in the 148-gene alignments were progressively removed in 4,000 position increments. The filtered alignments were individually subjected to rapid ML bootstrap analyses using RAxML. For each data point, we plotted the support values for (i) the sister relationship between strain PAP020 and fornicates (blue), (ii) the monophyly of fornicates (orange), (iii) the sister relationship between strain PAP020 and parabasalids (gray) and (iv) the monophyly of parabasalids (green).

**Figure S2:**
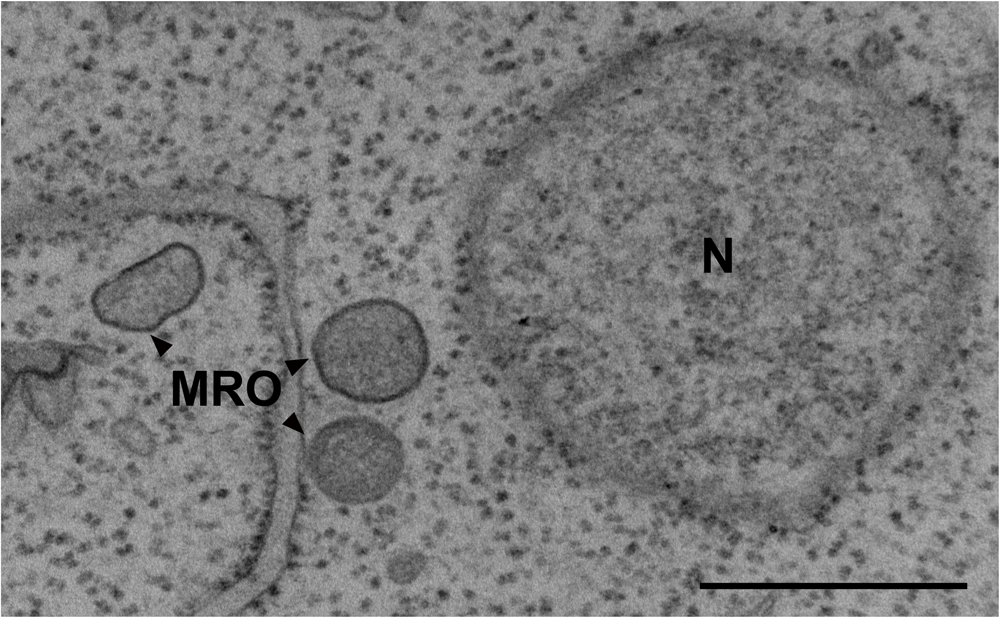
Transmission electron micrograph image of MRO of *Barthelona* sp. PAP020. Scale bar = 500 nm. A specimen for transmission electron microscopy (TEM) observation was prepared as follows; cultivated cells were centrifuged and fixed with pre-fixation for 1 h at room temperature with a mixture of 2% (v/v) glutaraldehyde, 0.1 M sucrose, and 0.1 M sodium cacodylate buffer (pH 7.2, SCB). Fixed cells were washed with 0.2 M SCB three times. Cells were post-fixed with 1% (v/v) OsO4 with 0.1 M SCB for 1 h at 4 °C. Cells were washed with 0.2 M SCB two times. Dehydration was performed using a graded series of 30–100% ethanol (v/v). After dehydration, cells were placed in a 1:1 mixture of 100% ethanol and acetone for 10 min and acetone for 10 min for two cycles. Resin replacement was performed by a 1:1 mixture of acetone and Agar Low Viscosity Resin R1078 (Agar Scientific Ltd, Stansted, England) for 30 min and resin for 2 h. Resin was polymerized by heating at 60 °C for 8 h. Ultrathin sections were prepared on a Reichert Ultracut S ultramicrotome (Leica, Vienna, Austria), double stained with 2% (w/v) uranyl acetate and lead citrate (Hanaichi et al., 1986, Sato, 1968), and observed using a Hitachi H-7650 electron microscope (Hitachi High-Technologies Corp., Tokyo, Japan) equipped with a Veleta TEM CCD camera (Olympus Soft Imaging System, Münster, Germany).

**Figure S3:**
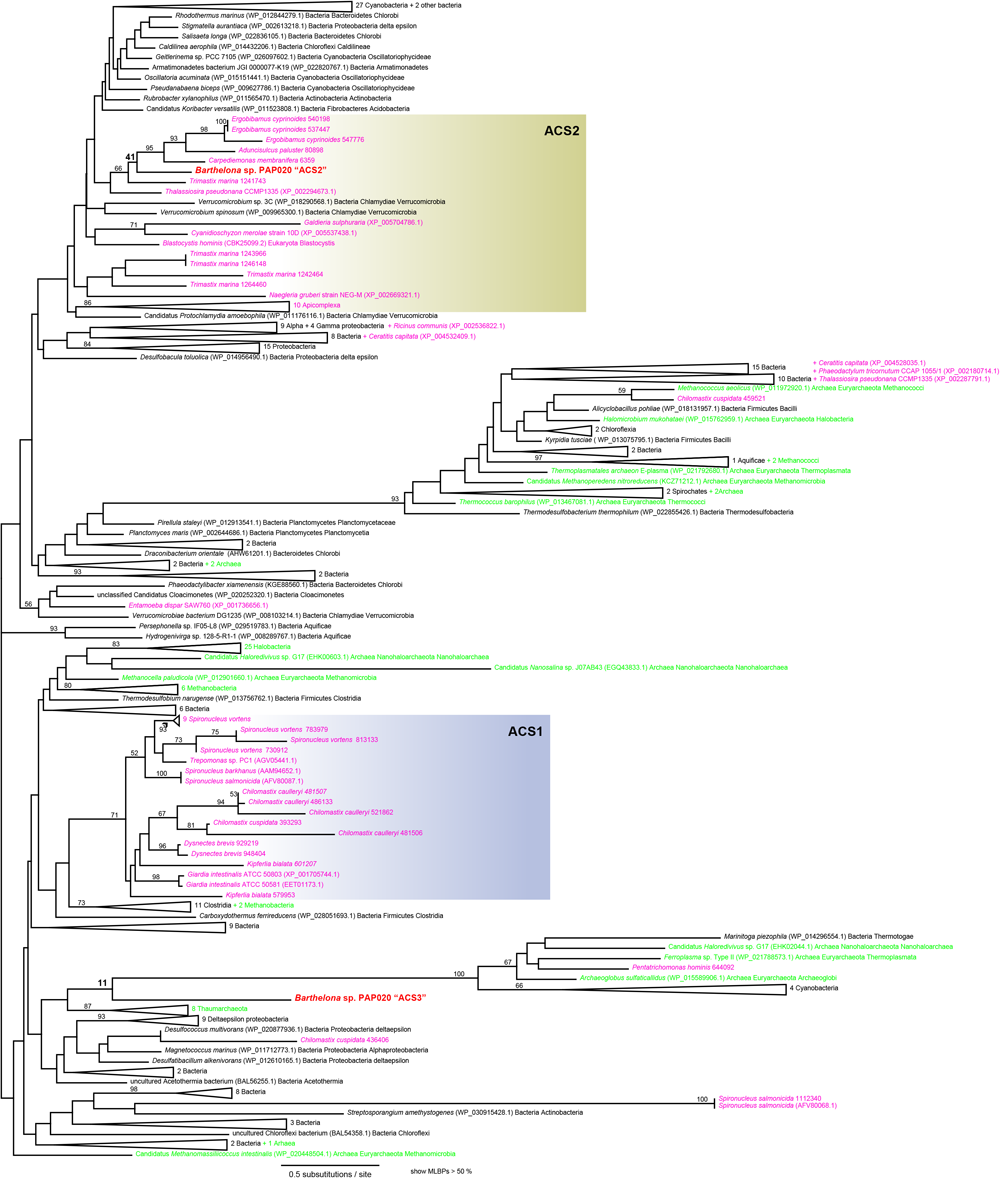
Phylogenetic tree of acetyl-CoA synthase (ACS) sequences. The ACS phylogeny was inferred using the maximum-likelihood (ML) method and ML bootstrap values (MLBPs) were mapped on the ML tree. MLBPs below 50% are not shown. Two ACS sequences of *Barthelona* sp. strain PAP020 are highlighted in red. The pink- and green-colored sequences are of eukaryotes and archaea, respectively. The clades of ACS1 and ACS2 were defined by referring to the phylogenetic analysis presented in Leger et al. (2017).

## References

Bernard C, Simpson AGB, Patterson DJ. 2000. Some free-living flagellates (protista) from anoxic habitats. Ophelia 52:113–142.

Bleidorn C. 2016. Third generation sequencing: Technology and its potential impact on evolutionary biodiversity research. Syst. Biodivers. 14:1–8.

Brown MW, Heiss AA, Kamikawa R, Inagaki Y, Yabuki A, Tice AK, Shiratori T, Ishida KI, Hashimoto T, Simpson AGB, et al. 2018. Phylogenomics places orphan protistan lineages in a novel eukaryotic super-group. Genome Biol. Evol. 10:427–433.

Burki F. 2014. The eukaryotic tree of life from a global phylogenomic perspective. Cold Spring Harb. Perspect. Biol. 6.

Burki F, Inagaki Y, Bråte J, Archibald JM, Keeling PJ, Cavalier-Smith T, Sakaguchi M, Hashimoto T, Horak A, Kumar S, et al. 2009. Large-scale phylogenomic analyses reveal that two enigmatic protist lineages, Telonemia and Centroheliozoa, are related to photosynthetic chromalveolates. Genome Biol. Evol. 1:231–238.

Burki F, Kaplan M, Tikhonenkov D V., Zlatogursky V, Minh BQ, Radaykina L V., Smirnov A, Mylnikov AP, Keeling PJ. 2016. Untangling the early diversification of eukaryotes: A phylogenomic study of the evolutionary origins of centrohelida, haptophyta and cryptista. Proc. R. Soc. B Biol. Sci. 283:20152802.

Burki F, Roger A, Brown MW, Simpson AGB. 2019. The new tree of eukaryotes. Trends Ecol. Evol. In press.

Cenci U, Sibbald SJ, Curtis BA, Kamikawa R, Eme L, Moog D, Henrissat B, Maréchal E, Chabi M, Djemiel C, et al. 2018. Nuclear genome sequence of the plastid-lacking cryptomonad *Goniomonas avonlea* provides insights into the evolution of secondary plastids. BMC Biol. 16:137.

Fukasawa Y, Tsuji J, Fu SC, Tomii K, Horton P, Imai K. 2015. MitoFates: Improved prediction of mitochondrial targeting sequences and their cleavage sites. Mol. Cell. Proteomics 14:1113–1126.

Gawryluk RMR, Tikhonenkov D V., Hehenberger E, Husnik F, Mylnikov AP, Keeling PJ. 2019. Non-photosynthetic predators are sister to red algae. Nature 572:240–243.

Grabherr MG, Haas BJ, Yassour M, Levin JZ, Thompson DA, Amit I, Adiconis X, Fan L, Raychowdhury R, Zeng Q, et al. 2011. Full-length transcriptome assembly from RNA-Seq data without a reference genome. Nat. Biotechnol. 29:644–652.

Haas BJ, Papanicolaou A, Yassour M, Grabherr M, Blood PD, Bowden J, Couger MB, Eccles D, Li B, Lieber M, et al. 2013. De novo transcript sequence reconstruction from RNA-seq using the Trinity platform for reference generation and analysis. Nat. Protoc. 8:1494–1512.

Janouškovec J, Tikhonenkov D V., Burki F, Howe AT, Rohwer FL, Mylnikov AP, Keeling PJ. 2017. A new lineage of eukaryotes illuminates early mitochondrial genome reduction. Curr. Biol. 27:3717–3724.e5.

Kamikawa R, Kolisko M, Nishimura Y, Yabuki A, Brown MW, Ishikawa S a, Ishida K, Roger AJ, Hashimoto T, Inagaki Y. 2014. Gene content evolution in Discobid mitochondria deduced from the phylogenetic position and complete mitochondrial genome of *Tsukubamonas globosa*. Genome Biol. Evol. 6:306–315.

Katoh K. 2002. MAFFT: a novel method for rapid multiple sequence alignment based on fast Fourier transform. Nucleic Acids Res. 30:3059–3066.

Katoh K, Standley DM. 2014. MAFFT: Iterative refinement and additional methods. Methods Mol. Biol. 1079:131–146.

Keeling PJ, Burki F. 2019. Progress towards the tree of eukaryotes. Curr. Biol. 29:R808–R817.

Kolisko M, Boscaro V, Burki F, Lynn DH, Keeling PJ. 2014. Single-cell transcriptomics for microbial eukaryotes. Curr. Biol. 24:R1081–R1082.

Kulda J, Nohýnková E, Cepicka I. 2017. Retortamonadida (with Notes on A *Carpediemonas-* like organisms and Caviomonadidae). In: Handbook of the Protists: Second Edition. Springer International Publishing. p. 1247–1278.

Kume K, Amagasa T, Hashimoto T, Kitagawa H. 2018. NommPred: Prediction of mitochondrial and mitochondrion-related organelle proteins of nonmodel organisms. Evol. Bioinformat. 14:1–12.

Lartillot N, Brinkmann H, Philippe H. 2007. Suppression of long-branch attraction artefacts in the animal phylogeny using a site-heterogeneous model. BMC Evol. Biol. 7:S4.

Lartillot N, Philippe H. 2004. A Bayesian mixture model for across-site heterogeneities in the amino-acid replacement process. Mol. Biol. Evol. 21:1095–1109.

Lartillot N, Philippe H. 2006. Computing Bayes factors using thermodynamic integration. Syst. Biol. 55:195–207.

Lax G, Eglit Y, Eme L, Bertrand EM, Roger AJ, Simpson AGB. 2018. Hemimastigophora is a novel supra-kingdom-level lineage of eukaryotes. Nature 564:410–414.

Lee WJ. 2002. Some free-living heterotrophic flagellates from marine sediments of Inchon and Ganghwa Island, Korea. Korean J. Biol. Sci. 6:125–143.

Lee WJ. 2006. Some free-living heterotrophic flagellates from marine sediments of tropical Australia. Ocean Sci. J. 41:75–95.

Leger MM, Kolisko M, Kamikawa R, Stairs CW, Kume K, Čepička I, Silberman JD, Andersson JO, Xu F, Yabuki A, et al. 2017. Organelles that illuminate the origins of *Trichomonas* hydrogenosomes and *Giardia* mitosomes. Nat. Ecol. Evol. 1:0092.

Li B, Dewey CN. 2011. RSEM: accurate transcript quantification from RNA-Seq data with or without a reference genome. BMC Bioinformatics 12:323.

Nguyen L-T, Schmidt HA, von Haeseler A, Minh BQ. 2015. IQ-TREE: A fast and effective stochastic algorithm for estimating maximum-likelihood phylogenies. Mol. Biol. Evol. 32:268–274.

Ronquist F, Teslenko M, Van Der Mark P, Ayres DL, Darling A, Höhna S, Larget B, Liu L, Suchard MA, Huelsenbeck JP. 2012. MrBayes 3.2: Efficient bayesian phylogenetic inference and model choice across a large model space. Syst. Biol. 61:539–542.

Shimodaira H. 2002. An approximately unbiased test of phylogenetic tree selection. Syst. Biol. 51:492–508.

Shimodaira H, Hasegawa M. 2001. CONSEL: for assessing the confidence of phylogenetic tree selection. Bioinformatics 17:1246–1247.

Simpson AGB. 2003. Cytoskeletal organization, phylogenetic affinities and systematics in the contentious taxon Excavata (Eukaryota). Int. J. Syst. Evol. Microbiol. 53:1759–1777.

Simpson AGB, Patterson DJ. 1999. The ultrastructure of *Carpediemonas membranifera* (Eukaryota) with reference to the “excavate hypothesis.” Eur. J. Protistol. 35:353–370.

Stamatakis A. 2014. RAxML version 8: A tool for phylogenetic analysis and post-analysis of large phylogenies. Bioinformatics 30:1312–1313.

Strassert JFH, Jamy M, Mylnikov AP, Tikhonenkov D V., Burki F. 2019. New phylogenomic analysis of the enigmatic phylum Telonemia further resolves the eukaryote tree of life. Mol. Biol. Evol. 36:757–765.

Susko E, Field C, Blouin C, Roger AJ. 2003. Estimation of rates-across-sites distributions in phylogenetic substitution models. Syst. Biol. 52:594–603.

Tanifuji G, Takabayashi S, Kume K, Takagi M, Nakayama T, Kamikawa R, Inagaki Y, Hashimoto T. 2018. The draft genome of *Kipferlia bialata* reveals reductive genome evolution in fornicate parasites. PLoS One 13: e0194487.

Tovar J, León-Avila G, Sánchez LB, Sutak R, Tachezy J, Van Der Giezen M, Hernández M, Müller M, Lucocq JM. 2003. Mitochondrial remnant organelles of *Giardia* function in iron-sulphur protein maturation. Nature 426:172–176.

Vincent AT, Derome N, Boyle B, Culley AI, Charette SJ. 2017. Next-generation sequencing (NGS) in the microbiological world: How to make the most of your money. J. Microbiol. Methods 138:60–71.

Wang HC, Minh BQ, Susko E, Roger AJ. 2018. Modeling site heterogeneity with posterior mean site frequency profiles accelerates accurate phylogenomic estimation. Syst. Biol. 67:216–235.

Yabuki A, Gyaltshen Y, Heiss AA, Fujikura K, Kim E. 2018. *Ophirina amphinema* n. gen., n. sp., a new deeply branching discobid with phylogenetic affinity to jakobids. Sci. Rep. 8.

Yabuki A, Kamikawa R, Ishikawa S a, Kolisko M, Kim E, Tanabe AS, Kume K, Ishida K-I, Inagaki Y. 2014. *Palpitomonas bilix* represents a basal cryptist lineage: insight into the character evolution in Cryptista. Sci. Rep. 4:4641.

Yubuki N, Inagaki Y, Nakayama T, Inouye I. 2007. Ultrastructure and ribosomal RNA phylogeny of the free-living heterotrophic flagellate *Dysnectes brevis* n. gen., n. sp., a new member of the Fornicata. J. Eukaryot. Microbiol. 54:191–200.

Yubuki N, Simpson AGB, Leander BS. 2013. Comprehensive ultrastructure of *Kipferlia bialata* provides evidence for character evolution within the Fornicata (Excavata). Protist 164:423–439.

Zhao S, Burki F, Bråte J, Keeling PJ, Klaveness D, Shalchian-Tabrizi K. 2012. *Collodictyon*-an ancient lineage in the tree of eukaryotes. Mol. Biol. Evol. 29:1557–1568.

